# Establishing a method for the cryopreservation of viable peripheral blood mononuclear cells in the International Space Station

**DOI:** 10.1101/2024.03.12.584560

**Authors:** Hiroto Ishii, Rin Endo, Sanae Hamanaka, Nobuyuki Hidaka, Maki Miyauchi, Naho Hagiwara, Takahisa Miyao, Tohru Yamamori, Tatsuya Aiba, Nobuko Akiyama, Taishin Akiyama

**Affiliations:** Laboratory for Immune Homeostasis, RIKEN Center of Integrative Medical Sciences, Yokohama 230-0045, Japan; Immunobiology, Graduate School of Medical Life Science, Yokohama City University, Yokohama 230-0045, Japan; Space Biomedical Research Group, Human Spaceflight Technology Directorate, JAXA, Tsukuba 305-8505, Japan; Japan Space Forum, Tokyo 101-0062, Japan

## Abstract

The analysis of cells frozen within the International Space Station (ISS) will provide crucial insights into the impact of the space environment on cellular functions and properties. The objective of this study was to develop a method for cryopreserving blood cells under the specific constraints of the ISS. In a ground experiment, mouse blood was directly mixed with a cryoprotectant and gradually frozen at -80 °C. Thawing the frozen blood sample resulted in the successful recovery of viable mononuclear cells when using a mixed solution of dimethylsulfoxide and hydroxyethyl starch as a cryoprotectant. Additionally, we developed new freezing cases to minimize storage space utilization within the ISS freezer. Finally, we confirmed the recovery of major mononuclear immune cell subsets from the cryopreserved blood cells through a high dimensional analysis of flow cytometric data using 13 cell surface markers. Consequently, this ground study lays the foundation for the cryopreservation of viable blood cells on the ISS, enabling their analysis upon return to Earth. The application of this method in ISS studies will contribute to understanding the impact of space environments on human cells. Moreover, this method may find application in the cryopreservation of blood cells in situations where research facilities are inadequate.

## Introduction

Throughout space missions, astronauts undergo an array of environmental changes that markedly differ from terrestrial conditions. These alterations encompass microgravity, exposure to high-energy space radiation, and psychological stress induced by confinement and fear, all of which may exert substantial influences on various physiological functions in human^1,2^. Several studies reported the impact of spaceflights on the immune systems^1,3,4^. For instance, various latent viruses were reportedly re-activated^5^. Moreover, analysis of blood samples from astronauts suggested that spaceflight influences the distribution of leukocytes^6,7^, natural killer cell function^8^, granulocyte and monocyte functions^7,9^, along with plasma cytokine levels^6,10^. Consequently, considering the implementation of short-term spaceflights for the public and long-term space exploration, including lunar and Mars missions, it is crucial to investigate the mechanisms through which spaceflights may affect immune function.

Single-cell RNA sequencing (scRNA-seq) technology has provided new insights into cell diversity and differentiation of immune cells^11^. In the NASA Twin Study^12,13^, scRNA-seq was employed to analyze the impact of spaceflight on gene expression and cell heterogeneity^13^. In this experiment, one twin served as the ground control and remained on Earth, while the other twin stayed aboard the ISS. For scRNA-seq, whole blood was collected into cell preparation tubes, and the microwell-based scRNA-seq analysis of peripheral blood mononuclear cells (PBMCs) was conducted. However, a limitation of this study is the absence of data collection during spaceflight; blood samples for scRNA-seq were obtained only before launch and after the return to Earth. Single-cell RNA-seq analysis of viable blood cells transported from the ISS may be compromised as the gene expression profile in these cells could be altered during the transportation to Earth. Whole blood cells in the cell preparation tube can be frozen for preservation, however, this freezing process induces cell lysis. Performing scRNA-seq analysis on a lysed blood cell mixture is extremely difficult, whereas bulk RNA-seq analysis of a mixed cell sample is feasible. Therefore, when generating scRNA-seq data from human PBMCs during spaceflight, it is crucial to preserve blood cells using a method that prevents their lysis^14^.

Cryopreservation is a widely-used method for generating viable cell stocks for long-term storage^15-17^. Density gradient centrifugation with Ficoll-Paque or cell preparation tubes is commonly employed for isolating PBMCs. Additionally, erythrocytes are eliminated by the addition of a buffer solution containing ammonium chloride, enhancing purity and suitability for downstream analysis. For cryopreservation, cells precipitated by centrifuge are resuspended in suitable cryoprotectants containing organic solvents such as dimethylsulfoxide (DMSO). While these methods are relatively straightforward in a laboratory setting, executing them in the ISS poses unique challenges. Due to the higher requirement of biosafety in the ISS as compared to the normal laboratory on Earth, blood and organic solvents need to be handled within a closed system. Consequently, opening tubes with human blood and organic solvents need to be avoided. Therefore, despite the feasibility of centrifugation, the task of removing the solution through aspiration and subsequently resuspending precipitated blood cells in cryoprotectants containing organic solvents presents considerable difficulties during spaceflights and missions, even though techniques to isolate and preserve cells under microgravity conditions were reported^18^. Consequently, these constraints in handling human blood and organic solvents could potentially hinder the cryopreservation of blood samples onboard the ISS.

The gradual cryopreservation of cells is crucial for minimizing intracellular ice crystal formation^15,16^, thereby reducing the probability of cellular demise. In addition to programmable cell freezers, various containers with optimized features for efficient cell freezing have been developed. Typically, these cell containers facilitate a gradual freezing process in cryoprotectant at a controlled rate of - 1 °C/min. These commercially available containers may impose significant spatial constraints within the ISS freezer and are also not applicable for cryopreservation using blood collection tubes. Consequently, it is important to establish a method for cryopreservation of blood cells within the ISS freezer without using a programmed cell freezer or commercially available containers.

We aim to establish a cryopreservation method suitable for execution aboard the ISS. In this study, we performed a ground experiment using murine blood samples and devices that can be used in the ISS environment. We found that freezing blood directly after adding a cryoprotectant containing dimethyl sulfoxide and hydroxyethyl starch, without additional additives like serum and albumin, allowed for the recovery of viable cells after thawing. Additionally, we report the development of a case designed for the cryopreservation of cells collected in a vacuum blood collection tube, which can be utilized in the ISS freezer. Overall, we propose that this procedure, along with a set of accompanying tools named the Biological sample MiXture device (BMX), would hold utility for the cryopreservation of viable blood mononuclear cells during missions on the ISS and thereby facilitate scRNA-seq and other analyses of PBMCs frozen aboard the ISS after their preservation and transport to Earth.

## Results

### Direct addition of cryoprotectant into EDTA-treated whole blood is applicable for cryopreservation of mononuclear cells

Due to the challenges of handling tubes containing both blood and organic solvents in the ISS environment, employing a standard cryopreservation method involving centrifugation, solution removal, and re-suspension in cryoprotectants becomes arduous. Therefore, we assessed a cryopreservation method involving the direct addition of cryoprotectants to an EDTA-treated blood sample^19^. We first tested whether two general cryoprotectants can be used in this method. Dimethylsulfoxide (DMSO) is known as a popular and convenient cryoprotectant^20^. In addition, previous studies indicated that the use of a cryoprotectant mixture containing hydroxyethyl starch and DMSO enhanced the recovery of unfractionated bone marrow cells from cryopreservation^21-23^. In the use of this cryoprotectants, the mixture of 12% hydroxyethyl starch and 10% DMSO (hereafter referred as to the CP-1 solution) with 8% human serum albumin (HSA) in saline is added to cell suspension solution in equal volumes ^21-23^. However, considering the richness of albumins in the mouse blood, we tested whether the addition of HSA could be omitted and replaced with phosphate-buffered saline. In all experiment conditions, equal volumes of cryoprotectant and mouse blood were mixed for the proper final concentration in a microcentrifuge tube. The tubes were enveloped with paper towels, a common bench-top method to facilitate a gradual cooling process without using cryogenic containers^17^, and then placed in a -80°C freezer. After being placed for 3 days at −80 °C, frozen cells were thawed and recovered. Upon visual inspection after thawing, substantial evidence of spontaneous hemolysis was apparent, suggesting that the freezing and thawing process caused blood cell degradation. In contrast, the survival of mononuclear cells was confirmed by staining with propidium iodide (counting for dead cells) and acridine orange (counting total cells with nucleus). Cryopreservation using the CP-1 solution yielded an average 75% survival ratio of mononuclear cells after the recovery and seems to be better than DMSO (average 57%) (Figure 1a). Notably, the survival ratio of recovered cells was above 60% for all trials using the CP-1 solution. In contrast, with DMSO, survival ratios in 3 trials among 5 trials were less than 50% (Figure 1a). Accordingly, within the framework of this cryopreservation methodology, the using the CP-1 solution appears to be superior to DMSO. The addition of HSA in the CP-1 solution may slightly improve cell survival (Figure 1a). Generally, it is recommended to mix HSA with the CP-1 solution immediately before adding them to cell suspensions. Therefore, considering the work efficiency in the ISS, we chose to use the CP-1 solution without HSA as the cryoprotectant in this method.

**Figure 1.**
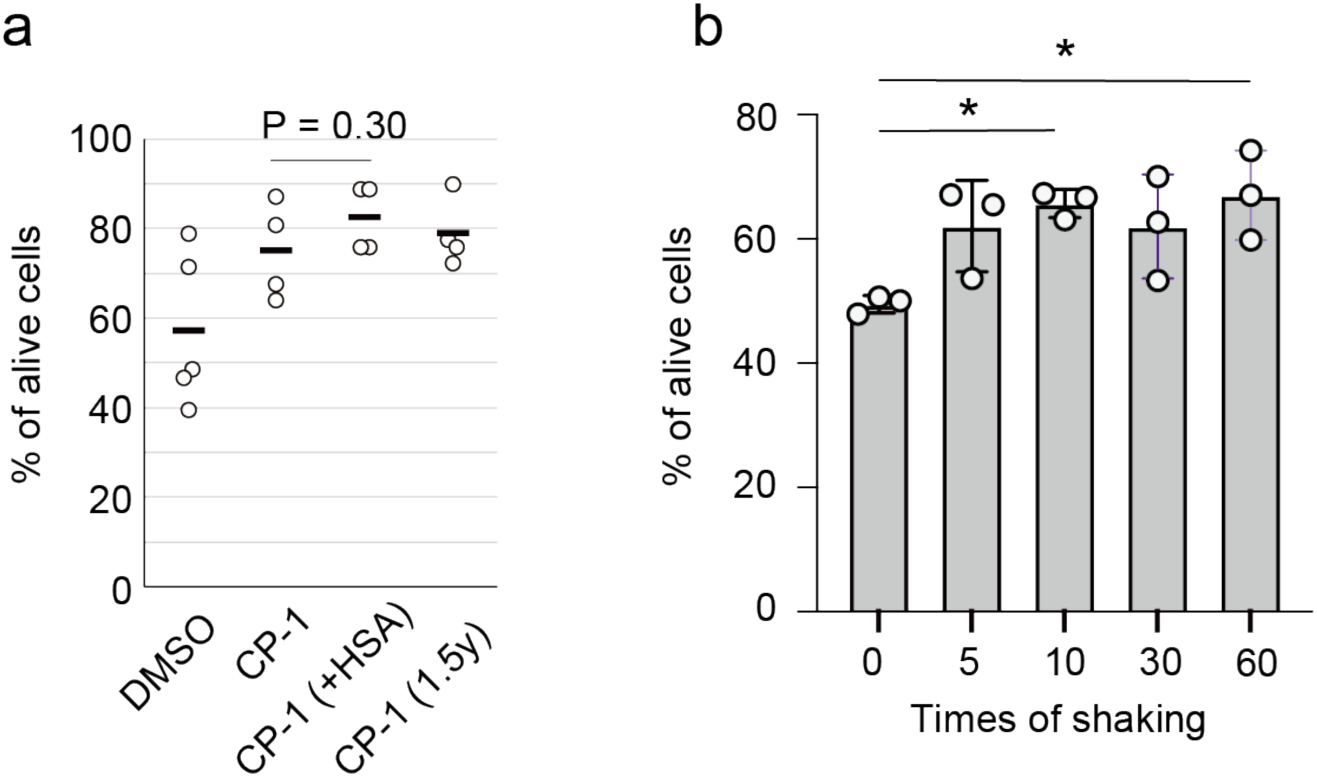
Cell survival ratio of recovered mononuclear cells in mouse blood. **a.** Comparison in cryoprotectants among DMSO, the mixed solution of DMSO and hydroxyethyl starch (CP-1) in the presence and absence of human serum albumin (HSA), and CP-1 stored in a syringe for 1.5 year at room temperature (1.5y). Samples (total 1 ml) were frozen in centrifuge tubes. N = 5 for DMSO, N = 4 for CP-1, N = 4 for CP-1 and HSA, and N = 2 for CP-1 for 6-month storage. Two-tailed and unpaired Student’s t-test. Bars indicate average values. **b.** Dependence of cell survival ratio on the mixing times after the cell recovery. After the addition of the cryoprotectant (CP-1) into the blood, the solution was mixed for the indicated time, wrapped with a paper towel, and frozen at −80 °C. After the recovery of the frozen cells, the viability was determined. Samples (total 4 ml) were frozen in blood collection tubes. N = 3. Average values are shown as boxes, and bars indicate standard deviation. *P < 0.05. One-way ANOVA and tukey’s test.

For the cryopreservation of blood during space missions, preserving cryoprotectants at ambient temperature for a long duration aboard the ISS would be preferable. We investigated the influence of long-term storage on the effectiveness of the CP-1 solution for cryopreservation. Following storage for 1.5 years in a plastic syringe at ambient temperature, the stored CP-1 solution underwent testing for blood cryopreservation. Data suggested that the storage for 1.5 years did not significantly influence the viability of recovered cells (Figure 1a), implying that the stored CP-1 solution can be used for cryopreservation in relatively long-term space missions.

Mingling two distinct solutions may pose challenges in the microgravity environment of the ISS. Additionally, astronauts might not always possess expertise in biological experiments involving cells and blood. Therefore, we investigated the optimal shaking frequency required to maintain maximum viability. We selected 7 ml vacuum blood collection tubes with 2 ml of fill volume to ensure sufficient free volume for adding 2 ml of CP-1 in the same volume. After collecting 2 ml of murine blood in each tube, 2 ml of the CP-1 solution was added to a 7 ml blood collection tube. Subsequently, collection tubes underwent mixing 0, 5, 10, 30, and 60 times with a shaking rate of 2 times/second (Figure 1b). To be slowly frozen, the tubes were enveloped in paper towels and placed in -80°C for several days. Notably, shaking 10 times proved sufficient, yielding an average of 60% survival upon recovery. However, in some instances, the survival ratio was slightly lower. Data deviated from the initial trial using microcentrifuge tubes (Figure 1a), likely due to an increase in total volume from 1 ml for the microcentrifuge tube to 4 ml for the blood collection tube. Survival rates consistently surpassed 50% with five or more shaking times, which would be deemed adequate for downstream analysis. However, in our following experiments, we shook the tubes 60 times to ensure sufficient mixing of the two solutions, considering the potential difficulty of mixing in microgravity conditions.

Our data suggest that directly adding the CP-1 solution as a cryoprotectant to the blood in the blood collection tubes is applicable for cryopreservation on the ISS. However, for biosafety reasons within the ISS laboratory, the possibility of splattering blood and organic solvents must be eliminated. Furthermore, the injection of the solution into a tube should be conducted safely and easily by astronauts. To address these concerns, we propose a procedure for injecting the CP-1 solution into the blood collection tube. As depicted in Figure 2, the CP-1 solution is loaded into a syringe before launch, and then, in the ISS, the pre-loaded CP-1 is injected using a safeguarded needle, called a blood transfer device.

**Figure 2.**
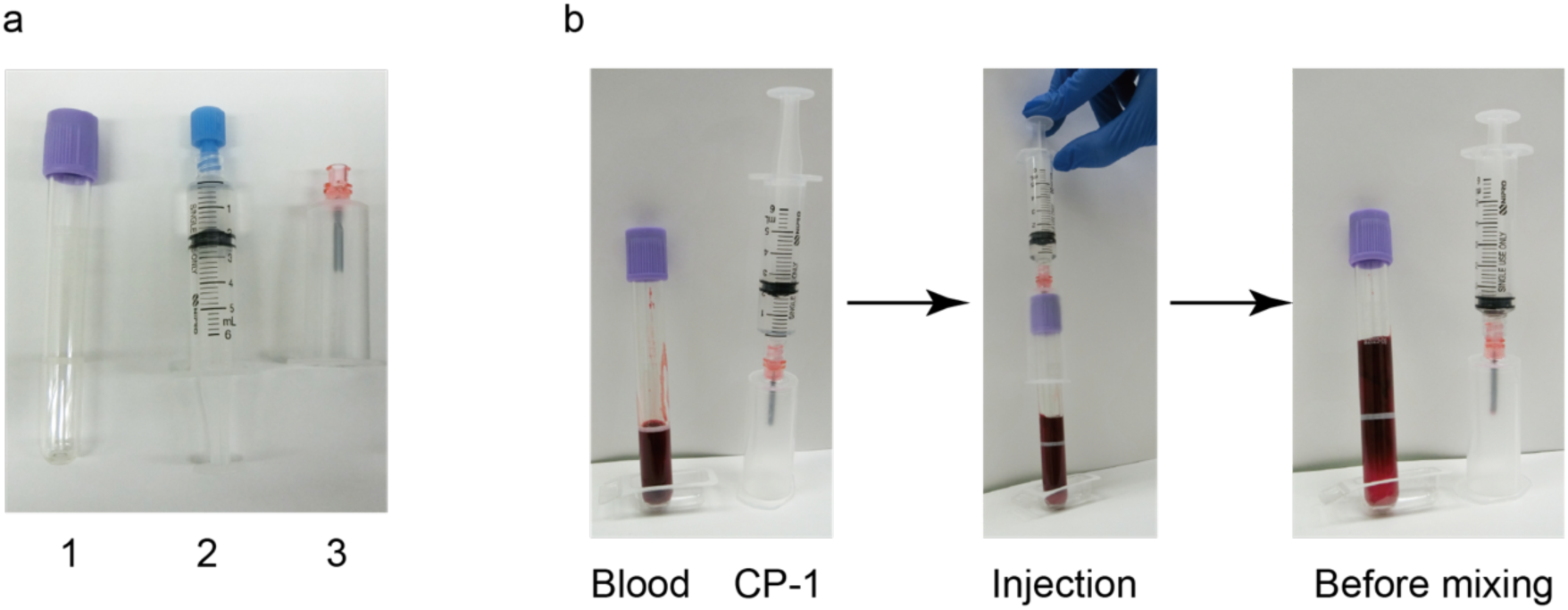
A typical procedure for the addition of the cryoprotectant into the blood collected in the collection tube. **a.** A set of materials for introducing CP-1 into blood within a collection tube includes: 1) the blood collection tube, 2) CP-1 in a syringe, and 3) a blood transfer device. **b.** A scheme for the procedure for adding CP-1 to blood in a collection tube. The CP-1 solution is pre-loaded into a syringe. After the blood collection in the tube, pre-loaded CP-1 is injected using a safeguarded needle (Injection). The needle is removed from the tube for shaking (Before mixing).

### The ethylene propylene diene monomer (EPDM) elastomer is suitable for a case designed for the slow freezing of blood samples

Although wrapping cryotubes with paper towels is a commonly used technique for slow freezing in cryopreservation^17^, it may be difficult for all astronauts to wrap tubes in the same way in the ISS. Moreover, commercially available containers do not accommodate the 7 ml vacuum blood collection tube. Therefore, we sought to design a specialized case specifically tailored for the slow freezing of blood in a standard blood collection tube within the ISS. As the material for the case, we chose ethylene propylene diene monomer rubber (EPDM), which is generally used as an insulator for sealing doors in freezers and refrigerators^24^. EPDM elastomers are light and maintain flexibility and elasticity even at low temperatures. Moreover, they resist degradation over time. These properties make them suitable for the materials of customized cases for freezing vacuum blood collection tubes in the ISS freezer. We utilized a tube-type of EPDM elastomer (6 mm and 10 mm thickness) that fits the blood collection tube (inner diameter, 16 mm) as the case for slow-freezing. We then assessed the feasibility of utilizing the EPDM elastomer tube as a slow-freezing case. Thus, we conducted a comparative study on the cooling rates of a solution (the 50% CP-1 solution) in blood collection tubes under 4 conditions: the tube placed in an EPDM elastomer case (6 mm or 10 mm thickness), the tube wrapped in paper towels, and the tube without any case or wrap. After being placed in the freezer (Figure 3a), the solution reached the zone of maximum ice crystal formation within 20 min and initiated the freezing process (Figure 3b). The cooling speeds of the steepest and most linear temperature change (around -10°C to -30°C) were -2.40°C/min for the 6 mm EPDM elastomer case, -2.20°C/min for the 10 mm EPDM elastomer case, and -2.05°C/min for the paper towel. The reduction rates of these cases were significantly slower than that of the unwrapped tube (- 5.01°C/min), and all conditions were practically comparable (Figure 3b and c). Consequently, data suggested that the EPDM elastomer cases can be used for the slow freezing of the solution in a blood collection tube.

**Figure 3.**
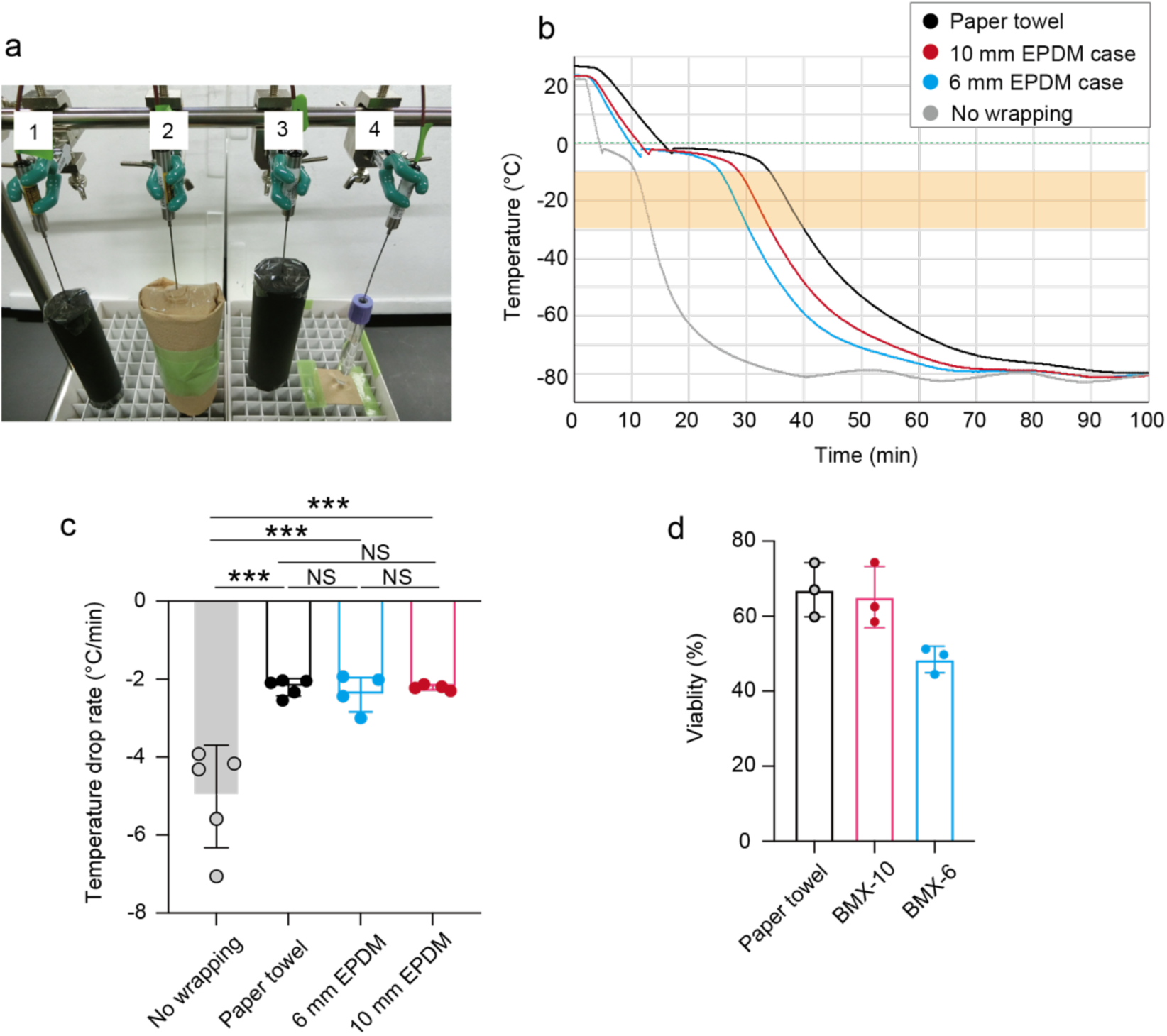
Development of an elastomer case used for slow cell freezing. **a.** A photograph for slow freezing of solutions using EPDM elastomer or paper towel wrapping in −80 °C freezer. Label 1, 6 mm EPDM elastomer case; label 2, paper towel; label 3, 10 mm EPDM elastomer case; label 4, without case and wrapping. **b.** A typical plot of decreasing rate in the temperature of the solution in tubes of the 6 mm EPDM elastomer case (blue), the 10 mm elastomer case (red), paper towels (black), or without wrapping (gray) at −80 °C. A green dot line indicates 0 °C. The orange shade indicates the temperature range exhibiting nearly linear cooling rates. **c.** Summary of cooling rates in the nearly linear temperature range. One-way ANOVA and tukey’s test. *** P < 0.001. NS indicates not significant. N = 5 for no wrapping, N = 5 for paper towel, N = 4 for 6 mm and 10 mm EPDM elastomer. Average values are shown as boxes, and bars indicate standard deviation. **d.** Cell survival ratio of recovered mononuclear cells in mouse blood using Biological sample MiXture device (BMX) and paper towel. N = 3 for each condition. Average values are shown as boxes, and bars indicate standard deviation.

### Cell-freezing by the direct CP-1 addition method causes a limited impact on the frequency of blood mononuclear cells

Our data suggested that the combination of the pre-loaded CP-1 solution in the syringe, blood transfer device, and collection tube, along with the EPDM elastomer case, may be used for the cryopreservation of blood in the ISS. This combination is named as the Biological sample MiXture device (hereafter referred as to BMX) with 6 mm (BMX-6) or 10 mm (BMX-10) thickness. We next tested that the BMXs are indeed appropriate for the recovery of viable blood mononuclear cells. Thus, the CP-1 solution was directly added to EDTA-treated mouse blood in the 7 ml blood collection tube. After mixing 60 times to complete the mixing, the tube was inserted into either case and placed in a −80 °C freezer. Frozen cells in the tubes were thawed after 3 days, and the cell survival ratio was determined (Figure 3d). BMX-10 demonstrated a cell viability of approximately 60%, comparable to the method using paper towels. In contrast, BMX-6 exhibited a slightly lower cell viability of around 50%. The data suggests that both BMXs enable the recovery of viable mononuclear cells from frozen blood, and can be used for various downstream analyses such as scRNA-seq.

We next investigated the influence of using the BMXs for cryopreservation on the frequency of cell populations in blood mononuclear cells, in comparison to the conventional method, which is cryopreserved by the CP-1 solution after eliminating red blood cells by the ammonium chloride solution (Figure 4). After the recovery, the use of BMXs caused a slight reduction in the total alive cell number per ml of blood as compared to the paper towel wrapping and the conventional method (Figure 4a). We next conducted a flow cytometric analysis on recovered cells, using 13 surface markers to identify major populations of mononuclear cells^25^ (Supplementary Figure 1). For flow cytometric data analysis, we employed a high-dimensional approach based on the expression levels of each marker. Specifically, we used UMAP dimensional reduction^26^ and k-means clustering^27^ to separate cell types (Figure 4b). The optimal number of clusters for the k-means algorithm was determined to be 10 using the elbow method^28^, which assists in identifying a suitable number of clusters based on the within-cluster sum of squares. We assigned these clusters based on the expression levels of surface markers (Figure 4c). Clusters 0 and 1, exhibiting high expressions of CD19 and MHC class II, were identified as B cells. Cluster 4, showing high expression of CD3e, was designated as T cells, while clusters 5 and 9, expressing high levels of NK1.1, were categorized as NK cells. B cell clusters were further separated by their expression levels of CD19 and MHC class II, which may reflect slight differences in the maturation stage. NK clusters were also divided into two groups, likely due to variations in the expression levels of NK1.1 and CD11b^29^. In addition to these lymphoid lineage cells, clusters 2, 3, 6, 7, and 8 were categorized as myeloid lineage cells based on expression of other markers. Cluster 8, with lower levels of CD45 and higher levels of FceR1, was identified as basophils. Cluster 6 was designated as eosinophils due to the expression of Siglec, while cluster 7 exhibited high expression of Ly6G, consistent with neutrophils. Clusters 2 and 3 were identified as monocytes, with cluster 2 characterized as Ly6C^lo^ and cluster 3 as Ly6C^hi^ monocytes^30^.

**Figure 4.**
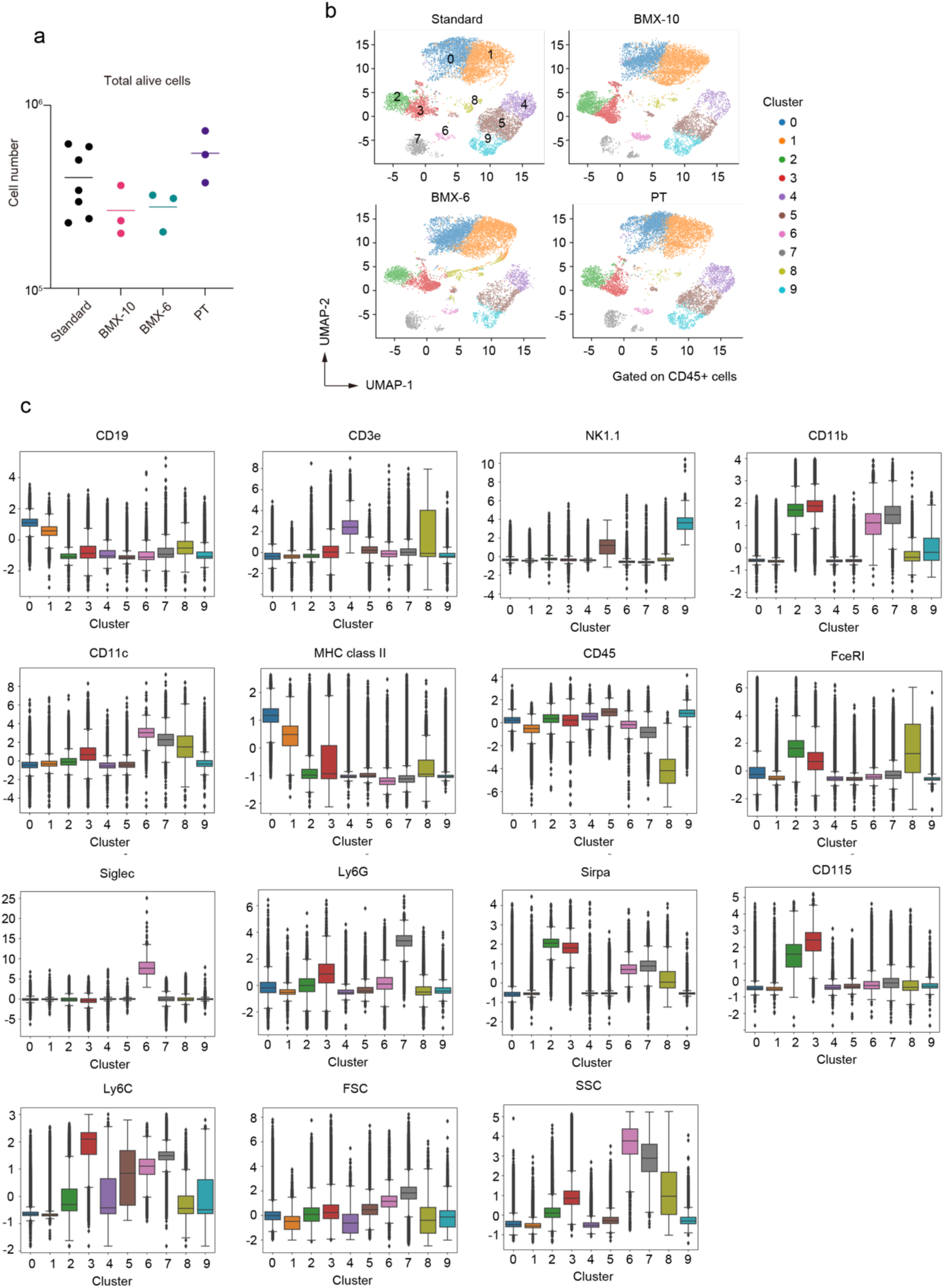
High-dimensional analysis of flow cytometric data. **a.** Recovered alive cell number after each cryopreservation method. Standard indicates the conventional method involving cryopreservation by the CP-1 solution after eliminating red blood cells by the ammonium chloride solution. Paper towels were used for wrapping tubes for the standard method. PT indicates the method of using paper towels instead of BMXs for wrapping tubes. N = 7 for standard, N = 3 for BMX-10, N = 3 for BMX-6, and N = 3 for PT. **b.** Uniform manifold approximation and production (UMAP) plot of flow cytometric data from mouse blood mononuclear cells after the recovery from cryopreservation. Cell clusters are separated by colors and numbers in the plot. All samples were integrated for the analysis, and dots exhibit 10,000 cells randomly selected from each data set. **c.** Expression level of surface markers in each cluster. The box is drawn from the first quartile to the third quartile. Bars indicate values of the minimum, the maximum, and the sample median. Outliers are shown as dots.

We proceeded to compare the influence of cryopreservation using BMXs on the frequency of these cell subsets. The data suggested that all major subtypes of mononuclear cells in the blood were present in the samples under all cryopreservation conditions (Figure 5). However, the ratio of some subsets in CD45^+^ cells was slightly but significantly altered when BMXs were utilized. Reductions in the ratio of T cells for both BMXs, an increase in basophil for BMX-6, and increases in Ly6C^hi^ monocytes and eosinophils for the BMX-10 were observed. Despite these limitations, since all major subsets are successfully recovered, we propose that the direct addition of the CP-1 solution to the whole blood using BMX can be employed for freezing blood in the ISS.

**Figure 5.**
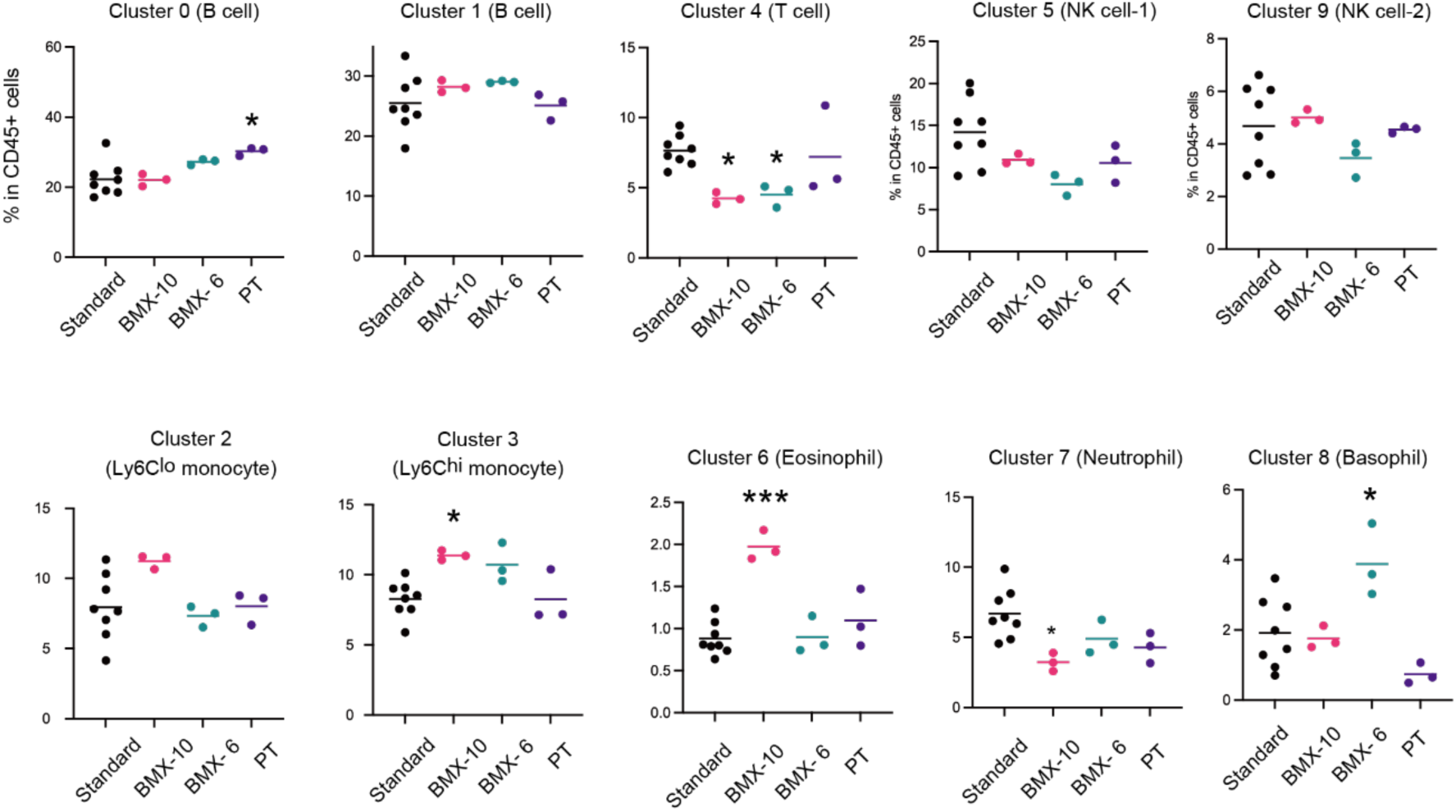
Comparison in the impact of cryopreservation methods on the ratios of cell clusters in recovered CD45^+^ mononuclear cells. * P <0.05, *** P < 0.001, indicating a significance in the difference from the standard. One-way ANOVA and tukey’s test. Labels are the same with that of Figure 4.

## Discussion

In this study, we developed a procedure wherein the CP-1 solution was directly added to EDTA-treated whole blood in a vacuum blood collection tube. During our investigation, Satpathy et al. reported the SENSE method, which involves freezing whole blood directly by adding a solution containing 80% heat-inactivated fetal bovine serum (FBS) and 20% DMSO at a 1:1 ratio^19^. In contrast to the SENSE method, our method does not use FBS but rather hydroxyethyl starch, which is widely employed as an artificial colloid plasma volume expander^31^. Given the extended storage of pre-loaded cryoprotectants at room temperature during the ISS mission, using a solution containing FBS in the ISS may not be feasible. We confirmed that the prolonged storage (1.5 year) of CP-1 at ambient temperature does not affect its efficacy as a cryoprotectant. Consequently, BMXs appear to hold promise as tools for cryopreserving blood on the ISS.

In microgravity, liquids exhibit different behaviors compared to Earth: Liquids are held together by surface tension, and convection does not occur. These characteristics of liquids in microgravity may hinder the mixing of two distinct solutions aboard the ISS. In this study, we examined the shaking times required for mixing the CP-1 solution with whole blood in 7 ml blood collection tubes. Our data suggest that 10 shaking cycles may be enough to mix them in this tube. However, given the specific behaviors of liquids in microgravity, we propose using 60 shaking cycles for mixing the CP-1 solution and whole blood in 7 ml vacuum blood collection tubes in the ISS. Notably, the 60-time shaking did not reduce the cell viability of recovered cells, suggesting this shaking speed (2 times/second) did not cause physical lethal damage to blood mononuclear cells.

After thawing and retrieval of blood cells from cryopreservation, typically 60% of mononuclear cells survived. Analysis of flow cytometric data suggests that this cryopreservation procedure enables the successful recovery of the majority of lymphoid or myeloid lineage cell populations, similar to the normal freezing procedure, albeit with a little more pronounced impact on T cell frequency. This indicates that our method is suitable for convenient cryopreservation in the ISS, where specialized laboratory equipment is not fully accessible.

Data suggested that the use of BMXs resulted in a decrease in the frequency of T cells following their recovery from cryopreservation. However, the underlying cause of this reduction remains unidentified. Given the diverse subtypes of T cells, further investigation is warranted to elucidate whether the cryopreservation by BMXs selectively influences specific subsets of T cells. Moreover, it should be noted that murine blood was used in this study. It is important to confirm that BMXs are applied for cryopreservation of human blood in the future.

In addition to this procedure, we developed a new type of case for cryopreservation of blood in the 7 ml vacuum blood collection tube. Our data suggested that wrapping by paper towel yielded the highest recovery ratio of viable cells after the cryopreservation. However, the method of wrapping the tube in paper towels may vary from individual to individual. Notably, the EPDM elastomer case is simple to use and is unlikely to vary from person to person. The 10 mm thickness of the case may be preferred over the 6 mm thickness due to data indicating a slightly better survival ratio when using BMX-10. However, the difference between these two was relatively small; therefore, the selection of these two tubes may depend on the allowed space in a freezer for each mission. Consequently, we propose that both BMX-6 and BMX-10 would be appropriate for freezing viable human PBMCs in the ISS.

The developed cryopreservation method and BMXs may hold significant promise for diverse applications in closed and microgravity environments such as the ISS. For instance, in standard scRNA-seq analysis, it is essential to use viable cells or cells that have been freshly fixed. The fixation of PBMCs using a fixative solution, such as paraformaldehyde may be difficult under the specific constraints of the ISS. Moreover, given that all blood cells, including red blood cells, could be fixed, only PBMCs need to be sorted from a pool of fixed blood cells containing a large number of fixed erythrocytes, which would present an additional challenge. Therefore, to perform scRNA-seq analysis to determine the status of PBMCs in humans aboard the ISS, viable PBMCs should be cryopreserved and returned to Earth. The BMXs that we developed in this study, would be useful for such purpose. Moreover, by using BMX, biological fluid samples collected in tubes can be pretreated by adding some reagents and other substances in closed and microgravity environments such as the ISS without opening the tubes.

In addition to usage in space missions, BMXs may be applicable to terrestrial research settings, facilitating the cryopreservation of blood cells for other medical studies. This would offer a potential breakthrough in preserving blood samples when research facilities are limited or unavailable, proving beneficial for medical research and diagnostics in resource-constrained environments. Consequently, BMXs and the cryopreservation technique developed for the ISS not only advance space-based research but also have far-reaching applications in medical science, healthcare, and emergency scenarios, making it a versatile and impactful innovation with broad-reaching potential.

## Methods

### Mouse

All mice were handled according to Guidelines of the Institutional Animal Care and Use Committee of RIKEN, Yokohama Branch (2018-075).

### Antibodies and reagents

Antibodies and reagents employed included FITC-anti-mouse NK1.1 (BioLegend, clone PK136, Cat#108705), PerCP-eFluor710-anti-mouse CD172a (eBioscience, clone P84, Cat# 46-1721-80), Alexa Fluor 647-anti-mouse Siglec F (BD, clone E50-2440, Cat# 562680), Alexa Fluor 700-anti-mouse MHCII (eBioscience, clone M5/114.15.2, Cat# 107621), APC-eFluor780-anti-mouse CD3e (eBioscience, clone 145-2C11, Cat# 47-0031-80), eFluor450-anti-mouse CD19 (eBioscience, clone eBio1D3 (1D3), Cat# 48-0193-80), BV510-anti-mouse CD11c (BioLegend, clone N418, Cat# 117338), BV605-anti-mouse CD115 (BioLegend, clone AFS98, Cat# 135517), BV650-anti-mouse/human CD11b (BioLegend, clone M1/70, Cat# 101239), BV711-anti-mouse Ly6G (BioLegend, clone 1A8, Cat# 127643), BV785-anti-mouse Ly6C (BioLegend, clone HK1.4, Cat# 128041), PE-Cy7-anti-mouse FceR1 (eBioscience, clone 44986, Cat# 25-5898-82), and BUV395- anti-mouse CD45 (BD, clone 30-F11, Cat# 565967).

### EPDM elastomer case for cryopreservation

The EPDM case was constructed from Aeroflex tubes (Aeroflex Co., ltd., Thailand), with thickness options of 6 mm and 10 mm. The main body of the case was formed by cutting the tubes to heights of approximately 125 mm for 6 mm thickness and 135 mm for 10 mm thickness. The lid was crafted from the same material of matching thickness, shaped into circular pieces with a diameter of 16mm to fit the inner diameter of the tube. Once the lid was placed into the bottom of the main body, it was securely taped in place. Following the insertion of the blood collection tube into the main body, the top was sealed using the lid.

### Cryopreservation

The blood of the mice (2 ml) was collected in a 7 ml blood collection tube (NIPRO, cat#: OP-EK0205). Without removing the cap, an equal volume of cell cryoprotective agent (CP-1® High Grade, KYOKUTO PHARMACEUTICAL INDUSTRIAL CO., cat#: 27207) was injected into the blood collection tube. At this point, a blood transfer device (BD Biosciences, cat#: 364880) is attached to the syringe containing the 2 ml of cryoprotective agent. After injecting the cell cryoprotective agent into the blood collection tube, the blood transfer device was removed from the tube while pressing and holding the plunger down. The blood collection tube was kept horizontally, and the blood was mixed with the cell cryoprotective agent by shaking the tube at indicated times along its long axis two times per second. Immediately after mixing, the mixture was placed into a freezing case or enveloped with paper towels (Kimtowel: NIPPON PAPER CRECIA CO., LTD., JAPAN), and frozen in a -80℃ freezer. For the small scale cryopreservation, 500 μl of the CP-1 solution (12% hydroxyethyl starch and 10% DMSO in saline) or 20% DMSO was added and mixed with 500 μl of murine blood in a 1.5 ml centrifuge tube. The tube was enveloped in paper towels and frozen in a -80°C freezer. For paper towel wrapping, a blood collection tube was wrapped in two paper towels and became approximately 4 cm in diameter after wrapping.

### Measurement of freezing rate

CP-1 (2 ml) and the same amount of saline solution (finally 50% CP-1 solution) were filled in a 7 ml blood collection tube at room temperature, and stored at a -80℃ freezer (Thermo Fisher Scientific Inc, MA) for around four hours. The temperature of the solution in blood collection tubes was measured at five-second intervals with Sheathed Thermocouples (CHINO CORPORATION, Tokyo JAPAN) and a data logger (MadgeTech, Inc., NH), under 4 conditions: the tube was placed in an EPDM elastomer case (6 mm or 10 mm thickness: Aeroflex Co., ltd., Thailand), the tube wrapped in paper towels (Kimtowel: NIPPON PAPER CRECIA CO., LTD., JAPAN), and the tube without any case or wrap.

### Recovery of cryopreserved cells

Frozen samples were thawed quickly by warming them in a 37°C water bath, with mixing every 30 seconds throughout the defrosting process. Two minutes after thawing began, half of the frozen sample started to defrost, at which point inversion mixing was repeated for 10 seconds. If the mixture was not sufficiently defrosted, an additional 5 seconds of mixing was provided to ensure thorough defrosting. Prior to complete defrosting, the blood sample was stored on ice. After defrosting, the blood sample was diluted with 40 mL of RPMI 1640 medium containing L-Glutamine and 5% FBS. Cell counts were performed after washing with a 2% FBS medium.

### Flow cytometric analysis

PBMCs were suspended in FACS buffer (2% fetal bovine serum; FBS in phosphate-buffered saline; PBS) containing 1 nM EDTA, and stained with antibodies in FACS buffer. 4’,6-diamidino-2-phenylindole (DAPI) was added to exclude dead cells. Data were acquired by using FACSymphony™ S6 Cell Sorter (BD Biosciences) and analyzed with FlowJo10. Machine learning was performed using PBMC data measured on a FACSymphony™ S6 Cell Sorter (BD Biosciences). In the PBMC dataset, live CD45^+^ cell data were extracted using FlowJo V10.6.2 (Treestar) and the subpopulation generation method implemented in flowkit^32^. The fluorescence intensity value of each stained antibody is obtained as a feature value because the measured data are flow cytometry data.

Therefore, the feature values were normalized respectively using the StandardScaler package implemented in sklearn. The dead cell marker and the features FSC-H/W and SSC-H/W were excluded from the obtained fluorescence intensities. Dimensional reduction was performed using UMAP^26^. Among the hyperparameters, n_neighbors (5, 25, 45, 65) and min_dist (0.1, 0.4, 0.7, 1.0) were tested under the 4 x 4 conditions and determined to be n_neighbors=25 and min_dist=0.4. K-means was used as the clustering method^27^, and the clustering was divided into ten populations based on the elbow method.

### Statistical analysis

A combination of One-way ANOVA and tukey’s test or two-tailed Student’s t-test was performed for statistical analysis.

## Data Availability

The datasets used during the current study are available from the corresponding author upon reasonable request.

## Author Contributions

**Hiroto Ishii**, Data curation, Formal analysis, Investigation, Validation, Writing; **Rin Endo,** Data curation, Formal analysis, Investigation, Validation, Writing – review and editing; **Sanae Hamanaka**, BMX investigation and development, Supervision, Writing; **Nobuyuki Hidaka**, BMX investigation and development; **Maki Miyauchi,** Investigation; **Naho Hagiwara,** Investigation; **Takahisa Miyao**, Supervision, Investigation; **Tohru Yamamori**, Supervision, Validation, Project administration; **Tatsuya Aiba**, BMX investigation and development, Supervision; **Nobuko Akiyama**, Supervision, Investigation, Writing – review and editing; **Taishin Akiyama**, Funding acquisition, Investigation, Project administration, Supervision, Validation, Writing.

## Conflict of Interest Statement

The authors declare no competing financial interests.

## Acknowledgment

This work is supported by JAXA feasibility study (TA) and CREST from the Japan Science and Technology Agency (JPMJCR2011) (TA).

## Figure and Figure legend

**Supplementary Figure 1.**
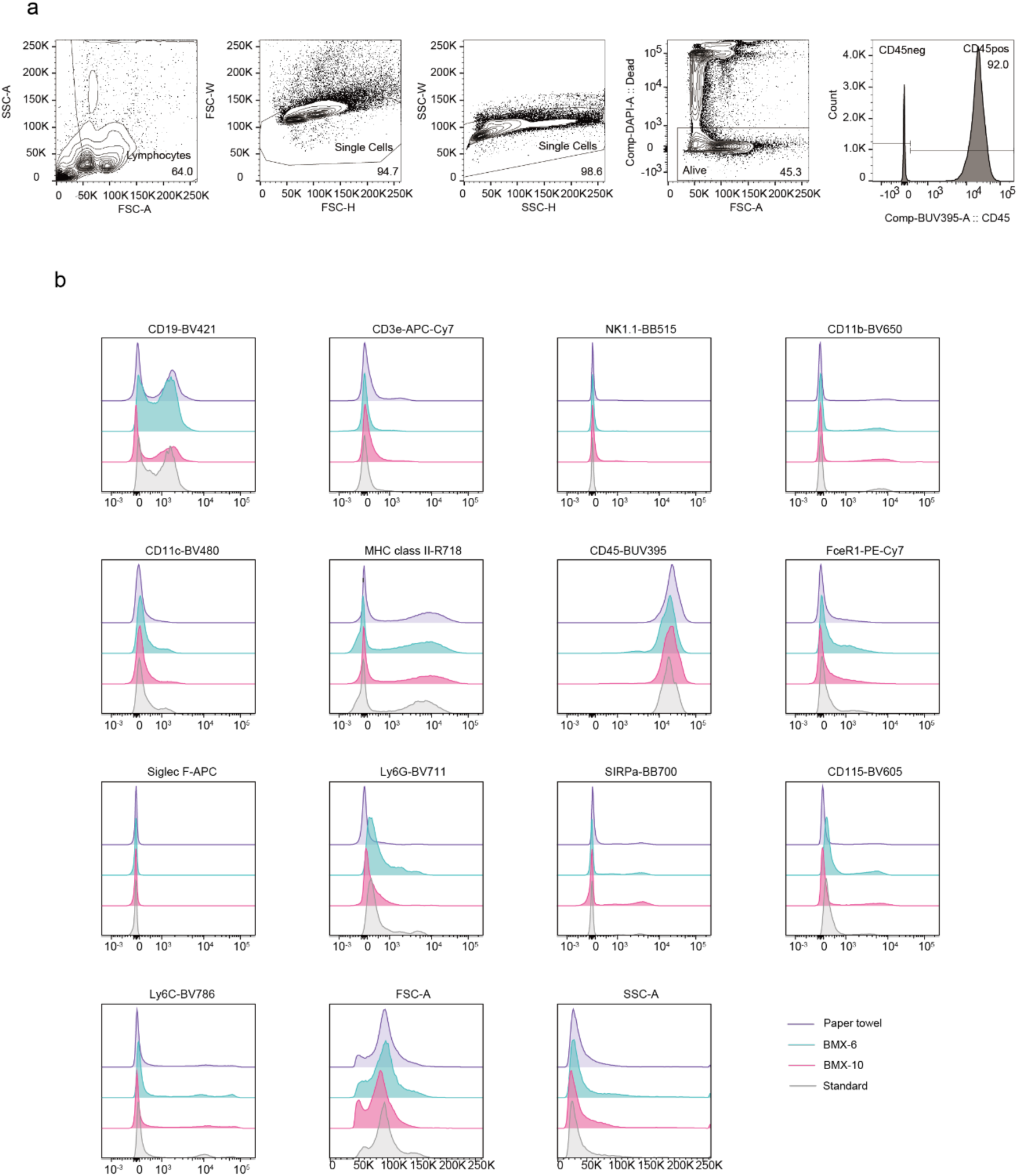
Gating strategy for flow cytometric analysis. **a.** Gating strategy for CD45^+^ cells. **b.** Flow cytometric profile of each cell surface marker. Typical data of each sample was exhibited. Purple histograms for wrapping by paper towel, blue histograms for BMX-6, red histograms for BMX-10, and gray histograms for the standard method involving the cryopreservation after the removal of red blood cells by hemolysis.

## Notes

### Competing Interest Statement

The authors have declared no competing interest.

